# Glutamatergic and GABAergic neuronal populations in the dorsolateral Periacqueductual Gray have different functional roles in fear conditioning

**DOI:** 10.1101/781187

**Authors:** Quentin Montardy, Zheng Zhou, Xuemei Liu, Zhuogui Lei, Pengyu Zeng, Chen Chen, Yuanming Liu, Kang Huang, Mengxia Wei, Liping Wang

## Abstract

It is though that only a subset of brain structures can encode emotional states. This can be investigated though a set of properties, including the ability of neurons to respond to a conditioned stimulus (CS) preceding an aversive unconditioned stimulus (US). The dorsolateral periacqueductal gray (dPAG) is a midbrain structure though to have an essential role in coordinating defensive behaviors in response to aversive stimulation. But its ability of dPAG neurons to encode a CS following fear conditioning as not been sufficiently studied.

Here we used calcium imaging by fiber photometry to record the activity of dPAG^VGluT2+^ and dPAG^GAD2+^ neuronal populations during unconditioned and conditioned aversive stimulation. Then, following an unconditioned stimulation we performed a retrieval experiment to quantify memory-like responses of dPAG neurons. This shown that whilst both dPAG^VGluT2+^ and dPAG^GAD2+^ neuronal populations respond to direct US stimulation, and to CS stimulation during conditioning, only the dPAG^VGluT2+^ population persisted in responding to the CS stimulation during retrieval. Finally, to better understand dPAG^VGluT2+^ and dPAG^GAD2+^ connectivity patterns, we performed a cell specific monosynaptic retrograde rabies virus tracing experiment. This revealed that different patterns of fibers projects to dPAG^VGluT2+^ and dPAG^GAD2+^, further complementing our recording showing divergences between PAG^VGluT2+^ and dPAG^GAD2+^ populations.

## Introduction

The dorsolateral Periacqueductual Gray (dPAG) is a midbrain structure known for its involvement in active defensive responses toward predators and pain [1] and is associated with autonomic responses [2]. The PAG is a nexus that receives highly processed information from areas that are involved in negative emotion, such as the amygdala and hypothalamus, and is thought to have an essential role in coordinating defensive behavior [1]–[3]. But the question of whether the dPAG is necessary for the emergence of fear states has rekindled debate over its role. Recently, Anderson and Adolphs proposed a set of properties to determine whether a brain structure can encode an emotional state [4]. Several of these have not been sufficiently studied in the dPAG; for example, the ability of dPAG neurons to respond to a conditioned stimulus (CS) preceding an aversive unconditioned stimulus (US) during fear conditioning, and the neural activity correlated with memory of the conditioning. To answer this question, we used calcium imaging by fiber photometry to record the activity of dPAG^VGluT2+^ and dPAG^GAD2+^ neuronal populations during unconditioned and conditioned aversive stimulation. Then, following an unconditioned stimulation we performed a retrieval experiment to quantify memory-like responses of dPAG neurons. Finally, to better understand dPAG^VGluT2+^ and dPAG^GAD2+^ connectivity patterns, we performed a cell specific monosynaptic retrograde rabies virus tracing experiment.

Here we shown that whilst both dPAG^VGluT2+^ and dPAG^GAD2+^ neuronal populations respond to direct US stimulation, and to CS stimulation during conditioning, only the dPAG^VGluT2+^ population persisted in responding to the CS stimulation during retrieval.

## Result

Here, we investigated PAG^VGluT2+^ neuronal population responses to aversive unconditioned and conditioned stimulation. Conditioning consisted of pairing a tone (CS+) with an aversive (footshock) unconditioned stimulus (US) (**Sup. Fig. 1.C**). In addition, a retrieval test was performed 24 h after conditioning to see whether PAG^VGluT2+^ neurons maintained a robust memory of the aversive conditioning. To record glutamatergic and GABAergic dPAG neuronal activity, we first injected a GCamp6s virus into the dPAG of VGluT2-cre and GAD2-cre mice, and implanted an optic fiber above the dPAG (**Fig. 1.A**). GCamp6s virus expression was mainly restricted to dPAG (occasionally overexpressing to dmPAG and lPAG) in VGluT2-cre animals (**Fig. 1.B**) and GAD2-cre animals (**Fig. 1.C**). Next, we delivered airpuff stimulation that evoked neuronal activity in dPAG^VGluT2+^ neurons, which required more than 9.9 s to return to baseline (**Fig. 1.D: red**), and evoked neural activity that returned to baseline around 4.7 s after stimulation in the dPAG^GAD2+^ neurons (**Fig. 1.D: blue**); this indicates that the VGluT2 and GAD2 dPAG populations may be associated with different temporal dynamics. On average, both glutamatergic and GABAergic dPAG neurons showed significantly higher responses compare to their respective baselines (VGluT2: BL=6×10^−4^dF vs. AP=5.2×10^−2^dF, P<0.0001; GAD2: BL=-1×10^−3^ dF vs AP=4×10^−3^ dF, P=0.0005; **Fig. 1.E**). Investigating trial-by-trial responses revealed that VGluT2 neuronal activity decreased gradually in a monotonic fashion (**Sup. Fig. 1.A**), whereas activity in the GAD2 population was more variable (**Sup. Fig. 1.B**). Next, the two groups were then subjected to aversive conditioning. As a control, mice received CS stimulation only one day before experiment. Neither dPAG^VGluT2+^ nor dPAG^GAD2+^ neurons responded to CS presentation alone (P>0.05, **Sup. Fig. 1.D;** P>0.05; **Sup. Fig. 1.G**). On the conditioning day, dPAG^VGluT2+^ neuronal activity significantly increased following CS-US pairing (BL=3×10^−3^ dF; vs CS=1×10^−2^ dF, P<0.05; vs US=4.8×10^−2^ dF, P<0.0001; **Fig. 1.G-H:** D1). Trial-by-trial analysis of neural responses following CS stimulation revealed an increase from baseline, but no change across trials (P >0.5; **Sup. Fig. 1.E-F**). US-evoked activity remained stable across trials (P < 0.01, **Sup. Fig. 1.E-F**). On the retention test day, VGluT2 neurons significantly increased following CS presentation (BL=-3×10^−4^ dF vs CS=1×10^−2^ dF, P<0.05; vs. Expected US=5×10^−3^, P>0.1; **Fig. 1.F-G**: d2). During conditioning, GAD2 neurons were significantly activated by CS and US stimulation (BL=-4×10^−4^ dF vs. CS=5×10^−4^ dF, P=1.5×10^−2^; vs. AP=4×10^−4^ dF, P<0.0001; **Fig. 1.H-I**: D1). Trial-by-trial analysis did not reveal any trend following CS-evoked responses; however, US-evoked responses were stable and increased significantly across trials (P <0.05; **Sup. Fig. 1.H-I**). Finally, GAD2 neurons did not respond to CS during the retention test (BL=-1.4×10^−4^ dF vs. CS=3.7×10^−4^ dF, P>0.1; **Fig. 1.H-I**: d2), and neither did they respond to expected-US (Expected US=2.8×10^−4^ dF dF, P>0.1).

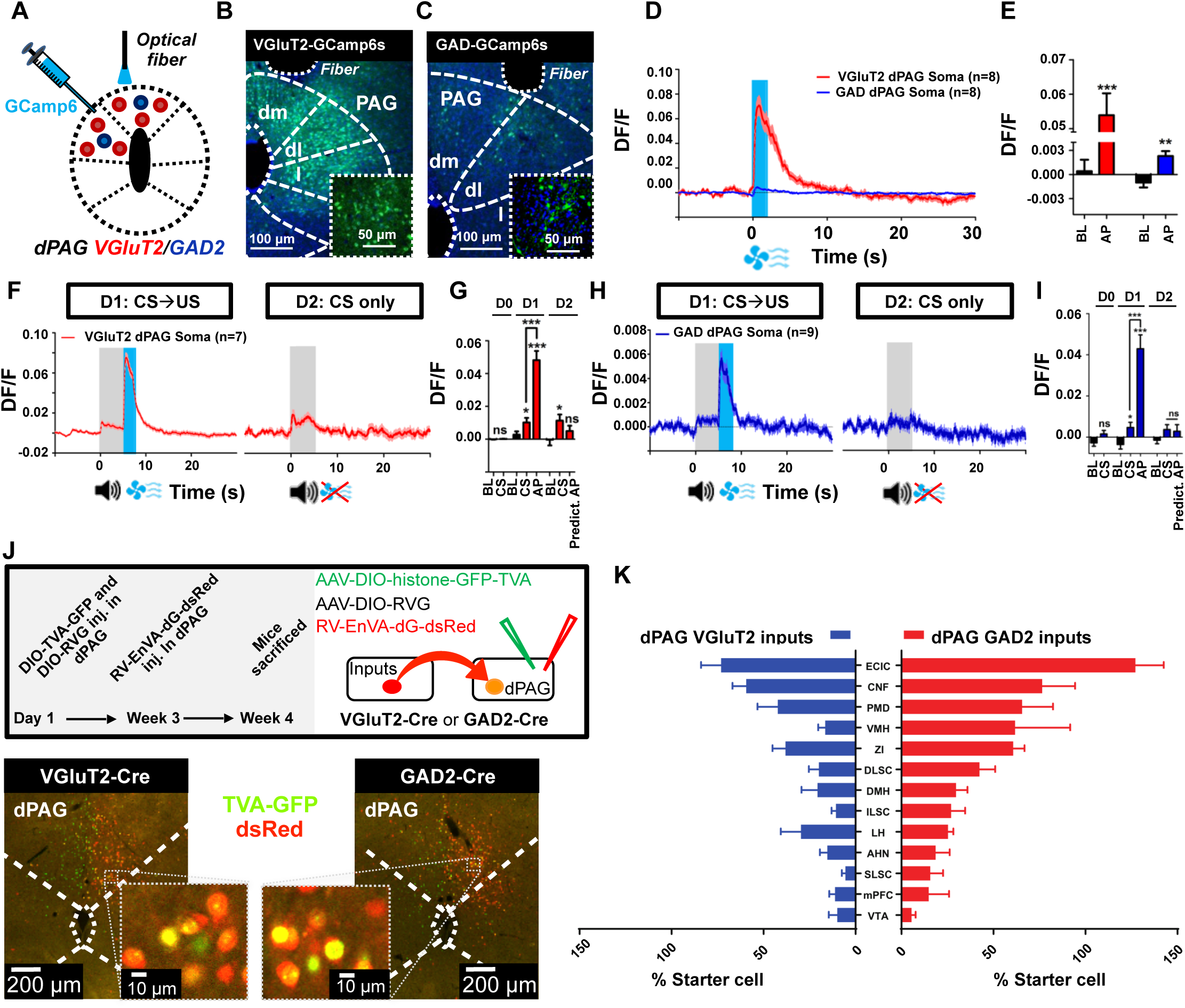
dPAG glutamatergic neurons still respond to CS during aversive retrieval test, while dPAG GABAergic neurons remain silent. **A.** AAV9-DIO-GCamp6s was injected in dPAG of VGluT2-ires-cre (red) or GAD2-ires-cre (blue), and an optical fiber was placed above dPAG. **B.** Representative image of GCaMP6s virus expression in dPAG of VGluT2, and **C.** GAD2 animals. **D.** Time course of signal in dPAG VGluT2 (red) and GAD2 (blue) neurons among all animals (VGluT2: n=8; GAD2: n=8) following airpuff-only. **E.** Averaged signal during baseline and airpuff stimulation. **F.** Averaged time courses for VGluT2::dPAG, which become sensitive to CS during D1 and remains sensitive in D2. **G.** Averaged Ca2+ signal during BL, CS, AP and expected AP, during D0, D1 and D2. **H.** GAD2::dPAG neurons are activated by CS during D1 but not D2. **I.** Averaged signal during CS, AP and expected AP at D0, D1 and D2. **J.** (Top) Schematic representation of RV-virus injection protocol. (Bottom) Representative picture of dPAG neurons expressing RV virus after injection in dPAG of VGluT2-cre animals (Red, rabies-dsRed; green, TVA-GFP; blue, DAPI, scale bar, 200 µm, and 10 µm representively). Boundaries of dPAG are drawn in white dashed lines. **K.** Percentage of starter cell projecting to dPAG VGluT2 neurons (in blue) or GAD2 neurons (in red)

Finally, to understand better how such differences occur between glutamatergic and GABAergic populations, we mapped their respective upstream projections and mapped the structures projecting to dPAG^VGluT2+^ and dPAG^GAD2+^ neurons using a Cre-dependent monosynaptic retrograde tracing technique. VGluT2-ires-Cre and GAD2-ires-Cre transgenic mice received AAV-CAG-DIO-histo-TVA-GFP (AAV2/9) and AAV-CAG-DIO-RG (AAV2/9) virus injections into dPAG. Three weeks later, dPAG was infected with RV-EvnA-DsRed (EnvA-pseudotyped, G-deleted and DsRed-expressing rabies virus) using the same coordinates. Mice were sacrificed one week after this second injection and injection sites were verified as dPAG (**Fig. 1.J**). Both neural populations receive projections from the same structures (**Fig. 1.K**). In particular, both populations receive afferent inputs from the external cortex of the inferior colliculus, the cuneiform nucleus, the dorsal part of the dorsal premammillary nucleus (PMD), and the zona incerta. However, while the GABAergic population receive strong inputs from the ventromedial hypothalamic nucleus (VMH), this is not the case for its glutamatergic counterpart.

In summary, both dPAG glutamatergic and GABAergic populations were sensitive to aversive stimulation. However, trial-by-trial analyses revealed different firing patterns between the two populations where only the glutamatergic population was responsive to CS during retention test, in a memory-like manner. By retrogradely characterizing afferences of these two populations, we revealed that they both receive upstream projections from the same structures, but with different strength of inputs, for example, the dPAG^GAD2+^ population receives many more VMH inputs than the dPAG^VGluT2+^ population does.

## Conclusion and Discussion

We recorded dPAG^VGluT2+^ and dPAG^GAD2+^ population activity using fiber photometry calcium imaging during unconditioned and conditioned aversive stimulation. Our main result is that, during a retrieval test after conditioning, only the dPAG^VGluT2+^ population persisted in responding to the CS stimulation. This indicates functional differences between glutamatergic and GABergic populations during the integration of aversive memories. These results are further supported by RV retrograde tracing data that show different patterns of fibers projecting to dPAG^VGluT2+^ and dPAG^GAD2+^.

First, we demonstrated that dPAG^VGluT2+^ and dPAG^GAD2+^ neuronal populations strongly respond to direct unconditioned aversive stimulation. Interestingly, the dPAG^VGluT2+^ population tended to gradually decrease its response across trials, although this was not statistically significant, whilst the dPAG^GAD2+^ population did not show any particular response pattern. These results suggest functional differences between glutamatergic and GABAergic neurons, and by extension also indicates that glutamatergic neurons, by maintaining a strong response for the CS 24 h after conditioning, may play a role to a form of memory. However, further experimentation is needed to systematically compare these two sub-populations, in particular, by integrating population responses uncovered by fiber optometry and single-unit recordings. In addition, it is important to note that we have only studied the response to aversive stimulation, and not for instinctive fear of predators [5] or other defensive behaviors [6]. It is quite possible that the neural responses of these neural populations may be different from each other in these specific contexts, but this is beyond the scope of this study.

Next, we submitted animals to an aversive conditioning session, to investigate the ability of dPAG neuronal population to encode signal value, as demonstrated by response to CS preceding an aversive US. We found that both dPAG^VGluT2+^ and dPAG^GAD2+^ populations strongly responded to CS following aversive conditioning, and in a similar manner. But in the retrieval session, during which the CS was delivered alone, only dPAG^VGluT2+^ and not dPAG^GAD2+^ population persisted in responding to the CS stimulation. First, this tends to confirm functional differences between the glutamatergic and GABAergic populations when it comes to formation of memory-like activity, as suggested by the unconditioned repeated stimulation. Then the fact that dPAG^VGluT2+^ population persisted in responding to CS retrieval is, on one hand, in accordance with a major part of the literature indicating a role for dPAG in memory and conditioning [7], [8]; whilst on the other hand, the absence of dPAG^GAD2+^ neuronal response can be viewed as in accordance with another study that shows a disconnect between emotion and behavior at the level of VMH to dPAG projections [9]. In this sense, our retrograde tracing data may reconcile previous reports in apparent contradiction: pathways originating from the VMH [9] or from other PAG columns [10], [11] may principally act on dPAG^GAD2+^ neurons and be involved in prolonged behavioral responses but not in memory; whilst parallel pathways originating from the VMH [7], [12] or the PMD [13] projecting to dPAG^VGluT2+^ neurons may participate in memory formation. This would suggest that glutamatergic neurons are the main contributors of aversive fear memory in dPAG, a population that could therefore contribute to the formation of fearful emotional states; whilst GABAergic neurons may partially belong to an independent dPAG circuitry that is only involved in the behavioral expression of fear, or in different categories of fear. Alternatively, the dPAG is also sending projections to structures such as the VTA [14]. Interestingly, VTA and dPAG glutamatergic neurons respond to aversive unconditioned and conditioned stimulation in a comparable manner [15]. Investigating dPAG to VTA function may help understanding the formation of aversive memory and emotion states.

Future experiments are required to investigate in detail whether dPAG^VGluT2+^ and dPAG^GAD2+^ neuronal populations are required for the acquisition and the expression of fear memory, and how they are intertwined at the microcircuit and circuit levels. In particular, investigating different fear categories may shed new light on the role of dPAG in the encoding of emotional states.

## Materials & Methods

### 1. Animals

All procedures were approved by Animal Care and Use Committees in the Shenzhen Institute of Advanced Technology (SIAT), Chinese Academy of Sciences (CAS). Adult (6-8 weeks old) male VGluT2-ires-cre (Jax No. 016963, Jackson Laboratory) and male Gad2-ires-cre (Jax No. 010802) transgenic mice were used in this study. All mice were maintained on a 12/12-h light/dark cycle at 25°C. Food and water were available ad libitum.

### 2. Viral preparation

For fiber photometry experiments, AAV2/9-EF1a-DIO-Gcamp6s virus was used. Virus titers were approximately 2-3×10^12^ vg/mL. For rabies tracing, viral vectors AAV2/9-EF1a-FLEX-TVA-GFP, AAV2/9-EF1a-Dio-histone-TVA-GFP, AAV2/9-EF1a-Dio-TVA-GFP, AAV2/9-EF1a-Dio-RV-G, and EnvA-RV-dG-dsRed were all packaged by BrainVTA Co., Ltd., Wuhan. Adeno-associated and rabies viruses were purified and concentrated to titers at approximately 3×1012 v.g/ml and 1×109 pfu/ml, respectively.

### 3. Viral injections

VGluT2-ires-cre and GAD2-ires-cre mice were anesthetized with pentobarbital (i.p., 80 mg/kg) and fixed on stereotaxic apparatus (RWD, Shenzhen, China). During the surgery, mice were kept anesthetized with isoflurane (1%) and placed on a heating pad to keep the body temperature at 35°C. A 10 µL microsyringe with a 33-Ga needle (Neuros; Hamilton, Reno, USA) was connected to a microliter syringe pump (UMP3/Micro4; WPI, USA) and used for virus injection into PAG (coordinates: AP:–3.8 mm, ML: –0.4 mm and DV:–2.45 mm).

### 4. Implantation of optical fibers

For photometry experiments, optical fibers (200 um in diameter, NA : 0.37) were chronically implanted in the PAG 3 weeks after virus expression. Optical fibers were unilaterally implanted above PAG (AP: –3.8 mm, ML: –0.5 mm and DV:–2.3 mm). After surgery all animals were allowed to recover at least 2 weeks to recover.

### 4. Conditioned airpuff test

We designed an apparatus consisting of a plexiglass tube (length 14 cm, diameter 10 cm) with a 1 cm groove for the optic fiber. The tube allowed horizontal movement and rotation. On the first day, mice were placed into the tube and received 5 trials of tone (80 db, 2000 Hz, 5 s) to habituate to the auditory stimuli. On the second day, 5 trials of conditioned stimuli (80 db, 2000 Hz, 5s) were presented to the animals after 3 min baseline, directly followed by 2 s unconditioned stimuli (air puff, 10-psi, 2 s). After 24 h, animals were place one last time in the apparatus and received 5 s conditioned stimuli without airpuff. Fiber photometry was recorded during the whole test.

### 5. Histology, immunohistochemistry, and microscopy

Mice were euthanized with overdose of chloral hydrate (300mg/kg, i.p) and transcardially perfused with ice-cold 1 x PBS and then with ice-cold 4% paraformaldehyde (PFA, sigma) in 1 x PBS. The brains were extracted from the skull and submerged in 4% PFA at 4°C overnight and then switched to 30% sucrose in 1 x PBS to equilibrate. Brains were cut into 40 µm thick coronal sections with cryostat microtome (Lecia CM1950, Germany). Freely floating sections were rinsed with PBS for 3 times and incubated in blocking solution (10% normal goat serum and 0.3% TritonX-100 in PBS) for 1h at room temperature. Then the slices were incubated for 24 h at 4°C with primary antibodies, anti-mouse TH (dilution 1:1000, M318, millipore) and anti-rabbit c-Fos (dilution 1:200, 2250, Cell Signaling). The secondary antibodies, Alexa fluor 488 anti-rabbit/goat (dilution 1:100, Abcam, ab150077) and Alexa fluor 405 anti-mouse/goat (dilution 1:100, Abcam, ab175660), were incubated at room temperature for 1 h. Finally, slices were mounted and cover-slipped in anti-fade aqueous mounting reagent with DAPI (ProLong Gold Antifade Reagent with DAPI, life technologies). The brain sections were imaged with Leica TCS SP5 laser scanning confocal microscope, and images acquisition was controlled by ImageJ software.

### 6. Fiber photometry

Ca^2+^ signals were recorded with a fiber photometry system (Thinker Tech, Nanjing). After AAV9-DIO-GCaMP6m virus injection, an optical fiber (NA: 0.37; NEWDOON, Hangzhou) was implanted into PAG as previously described. The photometry experiments were performed at least 10 days after surgery. To record Ca^2+^ signals, a laser beam (488 nm; Coherent, Inc. OBIS 488 LS) was reflected by a dichroic mirror (MD498; Thorlabs), focused on a microscope objective (Olympus, Inc., NA 0.37) and then coupled to an optical commutator (Doric Lenses). The laser intensity at the fiber tip was approximately 20 µW. Fluorescence was collected by a photomultiplier tube (PMT, R3896, Hamamatsu) and bandpass filtered by the dichroic filtered through a GFP bandpass emission filter (Thorlabs, Inc. Filter 510/30). The signals were amplified through a lock-in amplifier (Hamamatsu, Inc., C7319) and converted PMT current to voltage signal, which was further filtered through a low-pass filter (40 Hz cut-off; Brownlee 440). The analogue voltage signals were digitized and recorded by Cerebus electrophysiological recording system (BlackRock MicroSystem Inc.), and collected at a sampling frequency of 1000 Hz during the behavior test.

### 7. Photometry data analysis

Calcium Imaging signal was first extracted using Blackrock NPKM (Neural Processing Matlab Kit), according to bbbb instructions. Then analysis was carried on with Maltlab R2012a (The MathWorks Inc. ©) using custom code. Signal was analyzed as dF/F = (F - Fb)/Fb, where Fb is the baseline fluorescence before stimulation. Then data was smoothed using a 10 ms sliding windows. Time courses were made by averaging all individual trials aligned on the time of stimulation. Area surrounding time courses represent mean +/- SEM. Bar graphs represent all data averaged on the following periods: BL=[-2 s to 0 s before stimulation]; Airpuff-only AP and CS=[0 s to 3 s after stimulation]; D1 AP and D2 Predicted AP=[5 s to 7 s before stimulation]. Error bars represent mean +/- SEM. Trial-by-trial time courses and bar graphs were calculated using the same method, averaged across trial number. Significance was determined using two-tailed Student’s t-tests with a significance level of p<0.05.

## Acknowledgements

We thank Minmin Luo for providing us with GAD2-ires-Cre mice, Yangang Sun for VGluT2-ires-Cre mice.

This work was supported by National Natural Science Foundation of China (NSFC) 31630031 (L.W.), NSFC 31930047(L.W.), NSFC 81425010 (L.W.); International Partnership Program of Chinese Academy of Sciences 172644KYS820170004 (L.W.); Helmholtz-CAS Joint Research Grant GJHZ1508 (L.W.); the Strategic Priority Research Program of Chinese Academy of Science, XDB32030200; Guangdong Provincial Key Laboratory of Brain Connectome and Behavior 2017B030301017 (L.W.); JCYJ20170413164535041(L.W.), JCYJ20150401150223647 (Z.Z.); Shenzhen Municipal Funding GJHZ20160229200136090 (L.W.); Shenzhen Discipline Construction Project for Neurobiology DRCSM [2016]1379 (L.W.);Science and Technology Planning Project of Guangdong Province 2018B030331001(L.W.); CAS President’s International Fellowship 2017PB0090 (M.Q.); Guangdong Province International Scientific and Technological Cooperation 2019A050508008 (M.Q.).

## Author Contributions

M.Q., Z.Z., X.L. and Z.L. contributed equally to this work. M.Q. and Z.Z. designed and initiated the project. Z.Z. performed virus injections and fiber implantation. Z.L., and K.H. setup the behavior protocol. Z.L., P.Z., and C.C performed photometry experiments. Y.L., C.C. and K.H. helped to collect the data. M.Q. and M.W. processed and analyzed photometry data. X.L., Z.L. P.Z. and Y.L performed immunohistochemistry and quantitative analyzed of the tracing data. M.Q. and Z.Z. interpreted the results. M.Q, ZZ, X.L. and L.W commented the manuscript. M.Q., and L.W wrote the manuscript. L.W. supervised all aspects of the project.

---

**Supplementary figure 1:**
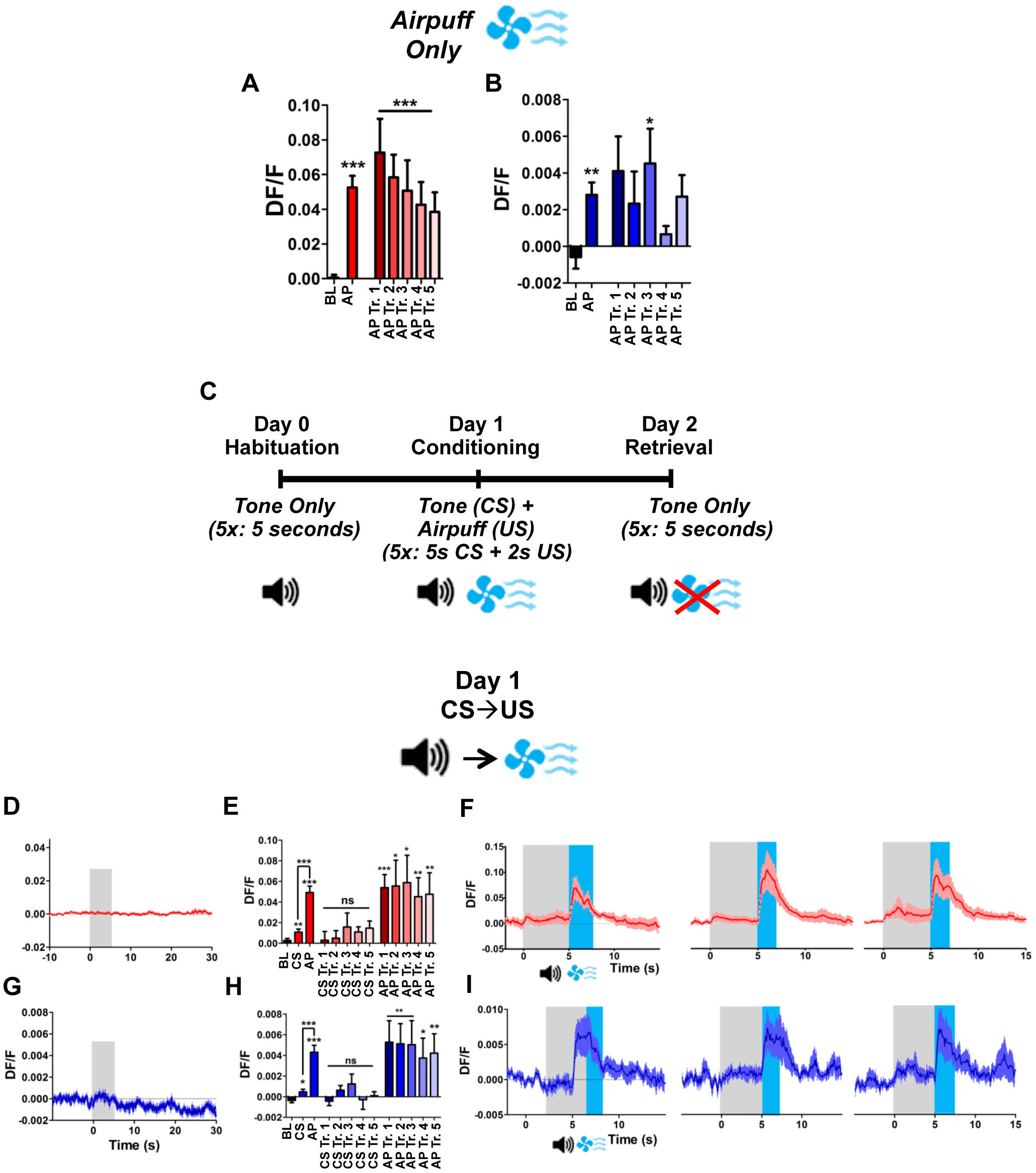
Trial by trial response of neurons during direct and conditioned aversive stimulation. **A.** Averaged trial by trial Ca2+ signal of VGluT2::dPAG animals during BL ([-2s:0s]) and Airpuff ([0s:2s]) in the Airpuff-only experiment. **B.** Averaged trial by trial Ca2+ signal of GAD2::dPAG animals. **C.** Schematic representation of CS-US conditioning, where CS is a 5 s tone and US a 2 s airpuff starting immediately after CS; Symbols represent tone CS and US airpuff time (respectively T=0s and T=5s). **D.** Averaged VGluT2::dPAG time course during D0 CS-only presentation. **E.** Averaged Ca2+ signal of VGluT2 dPAG neurons for the 5 first trials, averaged during BL ([-2s:0s]), CS ([0s:3s]), and AP ([5s:7s]) for conditioning experiment (D1). **F.** Time courses of the first (left) second (middle) and third trial (right) during conditioning experiment (D1), averaged among VGluT2::dPAG animals. **G.** Averaged GAD2::dPAG time course during D0 CS-only presentation. **H.** Averaged Ca2+ signal of GAD2 dPAG neurons for the 5 first trials, averaged during BL ([-2s:0s]), CS ([0s:3s]), and AP ([5s:7s]) for conditioning experiment (D1). **I.** Time courses of the first (left) second (middle) and third trial (right) during conditioning experiment (D1), averaged among GAD2::dPAG animals.

